# The human liver microenvironment shapes the homing and function of CD4^+^T-cell populations

**DOI:** 10.1101/2020.08.03.230953

**Authors:** Benjamin G. Wiggins, Laura J. Pallett, Xiaoyan Li, Scott P. Davies, Oliver E. Amin, Upkar S. Gill, Arzoo M. Patel, Konstantinos Aliazis, Yuxin S. Liu, Gary M. Reynolds, Gideon Hirschfield, Patrick T.F. Kennedy, Yuehua Huang, Mala K. Maini, Zania Stamataki

## Abstract

**Background & Aims:** Tissue-resident memory T cells (T_RM_) are important immune sentinels that provide efficient *in situ* immunity. Liver-resident CD8^+^ T_RM_ have been previously described, and contribute to viral control in persistent hepatotropic infections. However, little is known regarding liver CD4^+^ T_RM_ cells. Here we profiled resident and non-resident intrahepatic CD4^+^ T cell subsets, assessing their phenotype, function, differential generation requirements and roles in hepatotropic infection.

**Methods:** Liver tissue was obtained from 173 subjects with (n=109) or without (n=64) hepatic pathology. Multiparametric flow cytometry and immunofluorescence imaging examined T cell phenotype, functionality and location. Liver T cell function was determined after stimulation with anti-CD3/CD28 and PMA/Ionomycin. Co-cultures of blood-derived lymphocytes with hepatocyte cell lines, primary biliary epithelial cells, and precision-cut autologous liver slices were used to investigate the acquisition of liver-resident phenotypes.

**Results:** CD69 expression delineated two distinct subsets in the human liver. CD69^HI^ cells were identified as CD4^+^ T_RM_ due to exclusion from the circulation, a residency-associated phenotype (CXCR6^+^CD49a^+^S1PR1^-^PD-1^+^), restriction to specific liver niches, and ability to produce robust type-1 multifunctional cytokine responses. Conversely, CD69^INT^ were an activated T cell population also found in the peripheral circulation, with a distinct homing profile (CX_3_CR1^+^CXCR3^+^CXCR1^+^), and a bias towards IL-4 production. Frequencies of CD69^INT^ cells correlated with the degree of fibrosis in chronic hepatitis B virus infection. Interaction with hepatic epithelia was sufficient to generate CD69^INT^ cells, while additional signals from the liver microenvironment were required to generate liver-resident CD69^HI^ cells.

**Conclusions:** Intermediate and high CD69 expression demarcates two discrete intrahepatic CD4^+^ T cell subsets with distinct developmental and functional profiles.

**Graphical Abstract:** 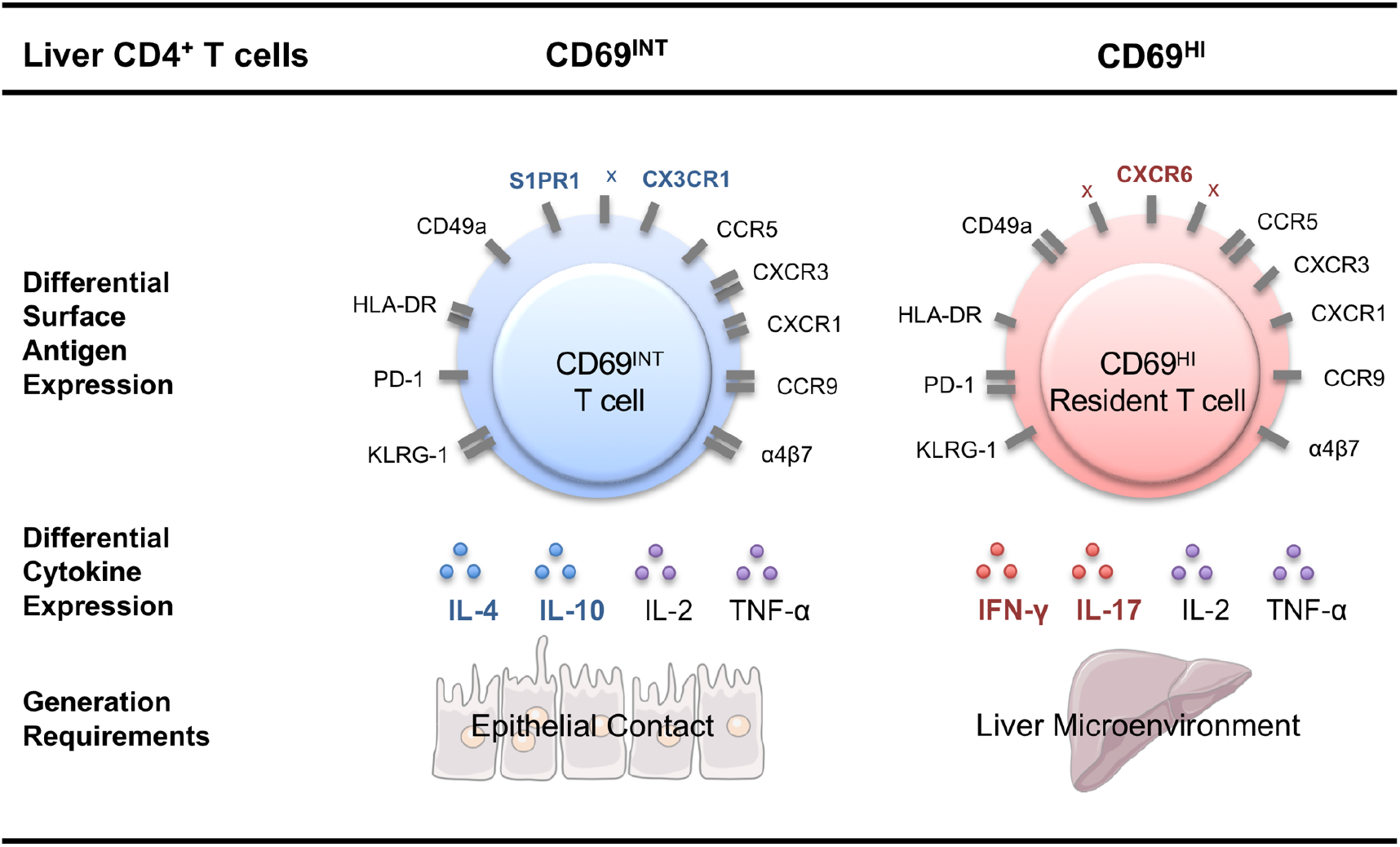

**Highlights:**

- CD69^HI^ (CXCR6^+^CD49a^+^S1PR1^-^PD-1^+^) are the CD4^+^ T_RM_ of the human liver
- Hepatic CD69^INT^CD4^+^ T-cells are distinct, activated, and recirculation-competent
- Stimulation evokes respective IFN-γ and IL-4 responses in CD69^HI^ and CD69^INT^ cells
- CD69^INT^ cell frequencies correlate with worsening fibrosis in chronic HBV patients
- Liver slice cultures allow differentiation of CD69^INT^ and CD69^HI^ cells from blood

**Lay summary:** Tissue-resident memory T cells (T_RM_) orchestrate regional immune responses, but much of the biology of liver-resident CD4^+^ T_RM_ remains unknown. We found high expression of cell-surface protein CD69 defined hepatic CD4^+^ T_RM_, while simultaneously uncovering a distinct novel recirculatory CD69^INT^ CD4^+^ T cell subset. Both subsets displayed unique immune receptor profiles, were functionally skewed towards type-1 and type-2 responses respectively, and had distinct generation requirements, highlighting the potential for differential roles in the immunopathology of chronic liver diseases.

## INTRODUCTION

Tissue-resident memory T-cells (T_RM_) are a nonrecirculating population that are critical in frontline adaptive immunity. Strategically positioned within tissues, these cells react to pathogen reexposure more efficiently than circulating memory subsets[1]. This function is mediated directly and by employing an innate-like ‘sensing and alarm’ strategy to enable recruitment and activation of other effector cells[2,3]. Human T_RM_ have now been identified in many organs[1, 4, 5], and differ substantially from their circulating counterparts in phenotype[6], function[7,8], metabolism[9, 10], maintenance[11] and responsiveness to stimuli[12]. Expression of tissue retention molecules CD69, CD103 and CD49a, and a lack of tissue egress markers including CCR7 and sphingosine-1-phosphate receptor 1 (S1PR1) define T_RM_[13]. Of these, CD69 is particularly important as a marker preserved on CD4^+^ and CD8^+^T_RM_ in all tissues[14], and separation through expression of this molecule alone has recently been used to define a human T_RM_ transcriptome with strong fidelity to more established murine T_RM_ profiles[13,15].

The liver is constantly exposed to dietary and bacterial products from the gut via the portal circulation, and has evolved as an important firewall to prevent pathogen dissemination[16]. Crucially the liver is considered immunotolerant, limiting responses to innocuous antigens[17]. However, pathogens, such as the hepatitis viruses, take advantage of this immunotolerant environment.

Recently, Pallett *et al*. identified intrahepatic CD8^+^T_RM_ (CD69^+^CD103^+^CXCR6^+^CXCR3^+^PD-1^+^ (PD-1 – programmed cell death protein-1)), capable of robust IL-2 production, associated with viral control in the liver of HBV-infected individuals[5]. However, little is known about CD4^+^T_RM_ and their role in liver defences. In one study, Wong *et al*. outlined distinct activation, differentiation and homing receptor profiles of liver perfusate CD4^+^T-cells as part of a multiorgan mapping study[18], supporting the possibility of a liver-resident CD4^+^T-cell population.

Here, we provide the first comprehensive phenotypic and functional analysis of intrahepatic CD4^+^T_RM_ in the human liver. We identified two distinct populations of CD69-expressing intrahepatic CD4^+^T-cells: CD69^HI^ and CD69^INT^. CD69^HI^CD4^+^T-cells within the human liver had prototypic hallmarks of tissue-residency including high expression of retention markers, exclusion from the circulation and rapid multifunctional type-1 cytokine production upon stimulation. We also report a novel population of intrahepatic CD69^INT^CD4^+^T-cells characterised by a unique chemokine receptor profile (CD69^INT^CX_3_CR1^+^CXCR3^+^CXCR1^+^). CD69^INT^ cells retained the ability to recirculate and upon stimulation produced the TH2 cytokine IL-4. Consistent with their TH2-type skewing, the frequency of CD69^INT^CD4^+^T-cells correlated with fibrosis progression in chronic hepatitis B. Finally, we demonstrated that contact with hepatic epithelia drives the CD69^INT^ phenotype, whilst CD69^HI^ cells required additional signals from the liver microenvironment.

## RESULTS

### CD69 expression on intrahepatic CD4^+^T-cells distinguishes three subsets with differential homing potentials

To identify intrahepatic CD4^+^T_RM_, we first analysed CD69 expression in 162 liver tissue samples from two research centres, the Centre for Liver and Gastrointestinal Research (University of Birmingham) and the Division of Infection and Immunity (University College London). Three intrahepatic CD4^+^T-cell phenotypes were identified: CD69^-^, CD69^INT^, and CD69^HI^ (**Fig.1A**). In order to test whether CD69^INT^ and CD69^HI^ cells represented distinct subsets, we assessed confinement to the liver, and expression of defined residency and homing signatures. While CD69^HI^CD4^+^T-cells were absent from the peripheral blood of all individuals tested, CD69^INT^ cells comprised up to 44% of the peripheral CD4^+^T-cell population (**Fig.1B**). Fine needle aspirate (FNA) sampling revealed that CD69^HI^ cells showed the strongest reduction in frequency compared to matched tissue biopsies, consistent with smaller fractions of interstitial T-cells sampled by FNA (**Supp.Fig.2A**)[21]. CD69^HI^ cells expressed high levels of tissue-retention molecules CD49a and CXCR6, and very little CX_3_CR1 and S1PR1, indicating an inability to egress into the circulation (**Fig.1C**). Furthermore, CD69^HI^ cells were enriched for an effector memory phenotype, a prerequisite for T_RM_ cells (**Supp.Fig.2B**). CD69^HI^ cells also contained the most CD103^+^ cells, but this percentage was low, as previously reported for CD4^+^T_RM_ (**Supp.Fig.2C**)[4].

**Fig.1.**
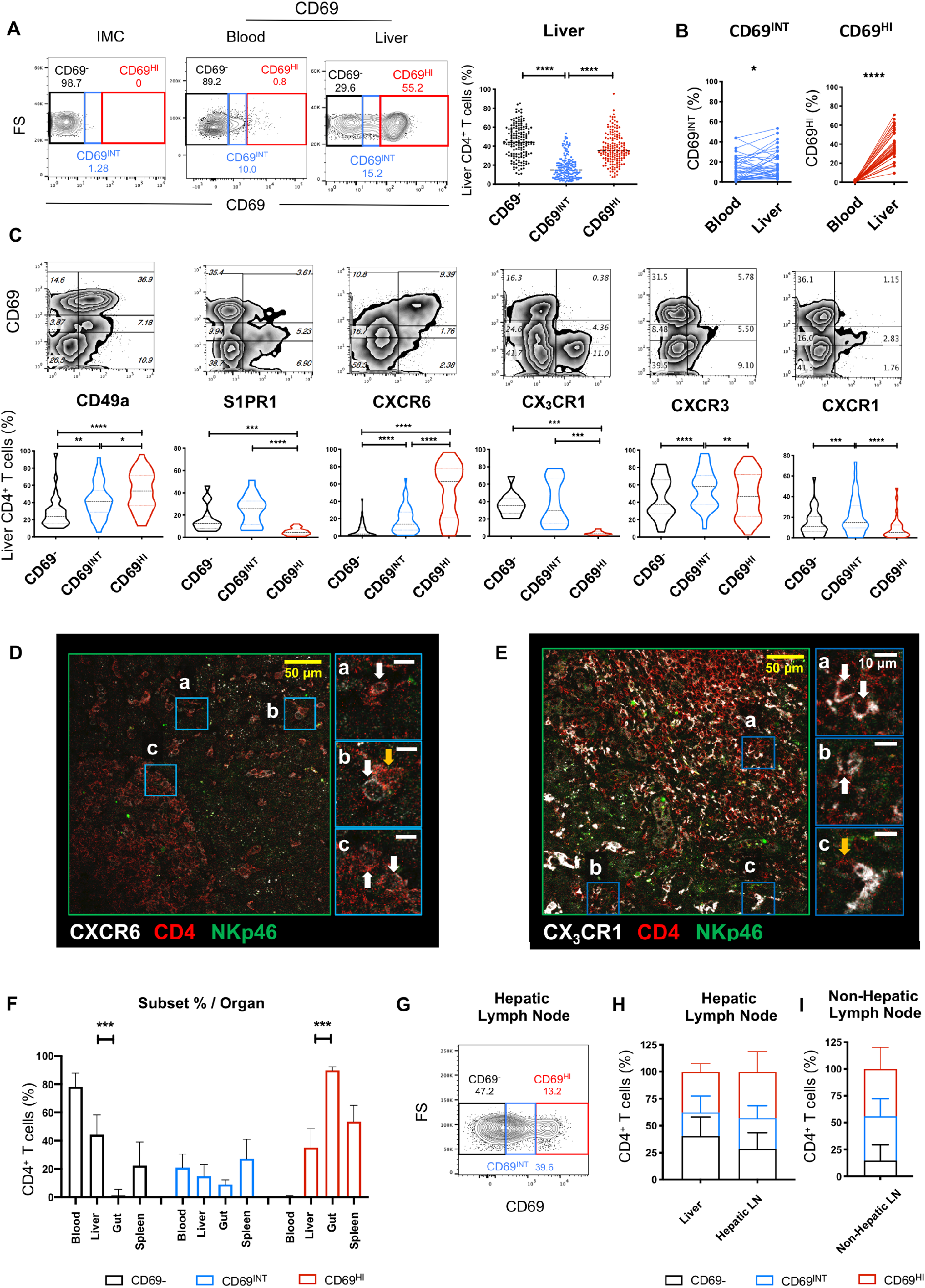
CD69 expression on intrahepatic CD4^+^T-cells distinguishes three subsets with differential homing potentials. **A** Gating strategy showing CD69^-^, CD69^INT^ and CD69^HI^ populations. Representative FACS plot for CD4^+^ subset distribution in blood and liver, and summary data showing % CD4^+^T-cells in IHL from two independent centres (n=162). **B** % CD69-expressing T-cell populations in paired blood and liver (n=39). **C** Expression of key homing and retention markers on CD69-expressing CD4^+^T-cell (from left to right, n=35, 16, 65, 12, 40, 40). Immunofluorescent staining of formalin-fixed liver sections showing localisation of **D** CD69^HI^ (NKp46^-^CD4^+^CXCR6^+^). Areas *a* and *b* mark sinusoids and *c* the outer portal lymphoid aggregate; and **E** NKp46^-^CD4^+^CX_3_CR1^+^ T-cells. Areas *a, b*, and *c* mark a portal lymphoid aggregate, liver parenchyma and a sinusoid respectively. White arrows indicate CD4^+^T-cells positive for indicated chemokine receptor whereas yellow arrows indicate chemokine receptor-negative CD4^+^T-cells. **F** CD69-expressing T-cell populations in blood (n=103), liver (n=162), gut (n=5) and spleen (n=3). **G** Representative CD69 expression on CD4^+^T-cells from a hepatic hilar lymph node (LN). **H** % of each subset in 6 matched liver and draining hepatic hilar LN samples, and (**I**) non-hepatic lymph nodes (n=5)

By contrast, hepatic CD69^INT^ cells expressed less CD49a, CXCR6 and CD103 than CD69^HI^ cells, although these residence markers were all expressed to a higher extent than on the CD69^-^ T-cells. Interestingly, CD69^INT^ cells retained expression of the tissue egress marker S1PR1 and fractalkine receptor CX_3_CR1, a chemokine receptor associated with migratory T-cells[13, 22, 23] (**Fig.1C**). CD69^INT^ were distinguishable from the other two populations by enhanced expression of chemokine receptors associated with parenchymal homing - CXCR3 and CXCR1[24]. Additional profiling revealed increased CCR5 and CCR9 expression on CD69^HI^ and CD69^INT^ respectively, and both CD69^+^ subsets expressed CCR6 than CD69^-^ CD4^+^T-cells (**Supp.Fig.2C)**. We also noted a reduction in cells expressing gut-associated α4β7 integrin expression for CD69^HI^ cells compared to other CD4^+^T-cells, and no differences in CCR10-expressing cells. These data show that the human intrahepatic CD4^+^T-cell compartment contains two distinct CD69-expressing subsets: a CD69^HI^ subset with *bona fide* T_RM_ features, and a CD69^INT^ subset with differential homing potential.

Next, we assessed whether differential expression of CXCR6 and CX_3_CR1 between the different intrahepatic CD4^+^T-cell populations affected their homing patterns. CXCR6^+^CD4^+^T-cells, enriched for CD69^HI^ cells, were found within sinusoids and the outer regions of portal lymphoid aggregates, similar to previous reports on CD8^+^T_RM_[5] (**Fig.1D**). By contrast, CX_3_CR1^+^CD4^+^T-cells were distributed throughout portal aggregates, parenchymal regions (**Fig.1E**), and in close proximity to large veins (**Supp.Fig.2D**). These data support the differential tissue distribution of CD69^INT^ and CD69^HI^ cells, suggesting a more discrete vascular sinusoidal/periportal niche for CD69^HI^ cells, whereas CD69^INT^CX_3_CR1^+^ T-cells could additionally access the liver parenchyma.

To address whether CD69^INT^CD4^+^T-cells were a liver-specific population, we examined the presence of all three subsets in samples obtained from the human gastrointestinal tract and spleen. CD69^INT^ were also present in these tissues, however CD69^HI^ cells represented a much greater proportion of all CD4^+^T-cells in the gut compared to the liver, and similar in the spleen, hepatic hilar lymph nodes and distal non-hepatic lymph nodes (**Fig.1F-I**). We reasoned that the expression of tissue-egress markers S1PR1 and CX_3_CR1 would imbue CD69^INT^ cells with the ability to recirculate through lymphatics[22]. In support, both hepatic hilar and distal non-hepatic lymph nodes contained comparable proportions of CD69^INT^ cells to the liver (**Fig.1G-I**). Combined, these data all point to CD69^HI^ cells encompassing CD4^+^T_RM_ in the liver, with CD69^INT^ cells a distinct subset implicated in wider tissue and lymphatic immunosurveillance.

### CD69^HI^ T_RM_ demonstrate a restrained, resting phenotype, while CD69^INT^ cells exhibit features of activation

Alongside a tissue-residence marker, CD69 has traditionally been used as a marker of early lymphocyte activation[25]. Therefore, we examined the activation status of the three hepatic CD4^+^ populations. Intriguingly the extent of cellular activation did not correlate with levels of CD69 expression in the liver; CD69^INT^ cells were the most activated subset with increased expression of CD38 and HLA-DR (% and MFI, **Fig.2A**). Consistent with recent activation *in vivo*, the CD69^INT^ population also showed higher proliferation than CD69^HI^ cells *ex vivo* with the highest percentage of Ki-67^+^ cells (**Fig.2B**). Regulatory T-cells (T_REG_, CD4^+^CD25^HI^CD127^LO^) were not significantly enriched in any CD4^+^ subset, and T_REG_ functional markers cytotoxic T-lymphocyte-associated protein-4 (CTLA-4) and CD39 were similarly expressed by both populations compared to CD69^-^CD4^+^T-cells (**Supp.Fig.3**).

**Fig.2.**
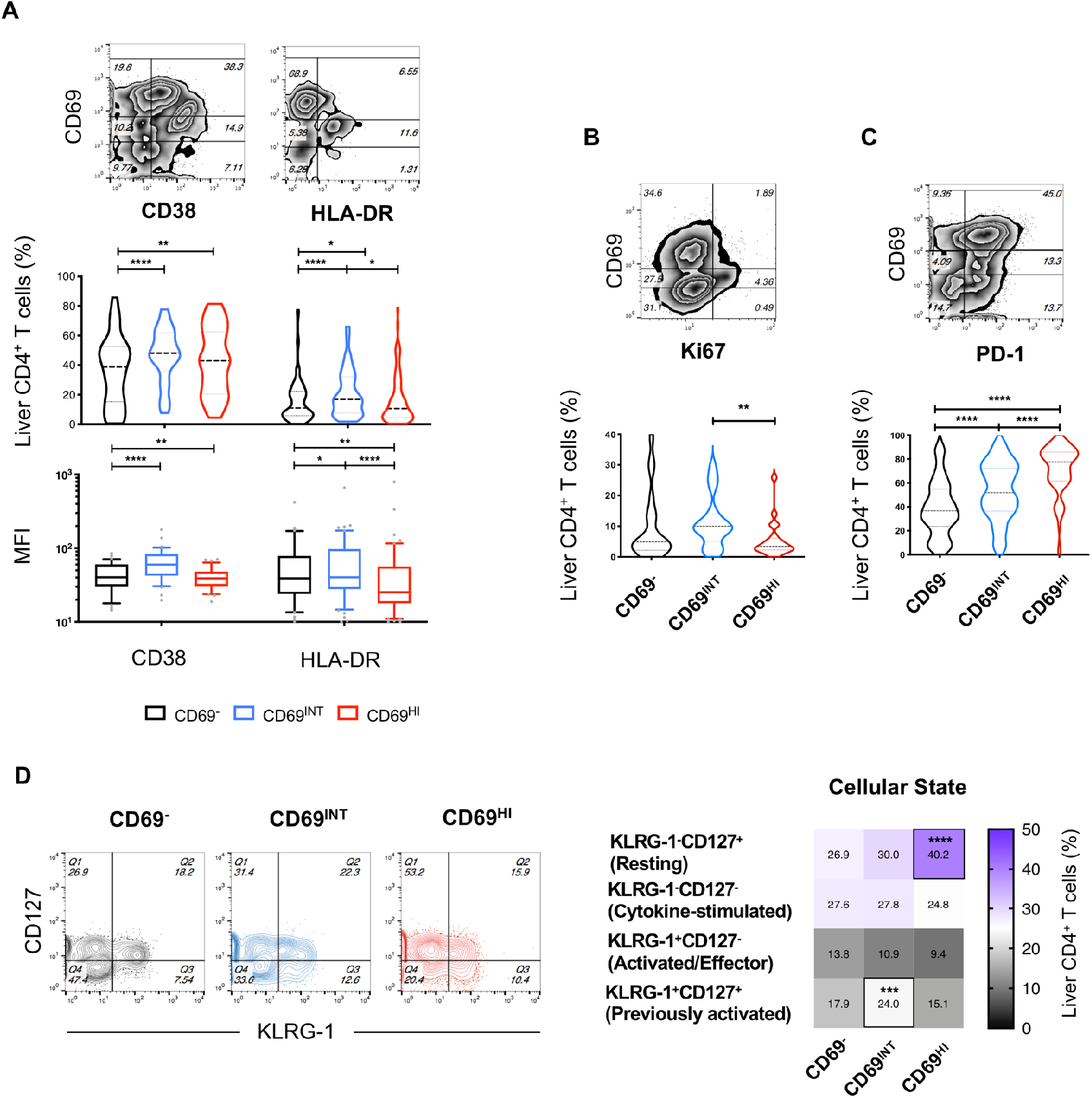
CD69^HI^T_RM_ demonstrate a restrained, resting phenotype, while CD69^INT^ cells exhibit features of activation. **A** % and mean fluorescence intensity (MFI) expression of CD38 (n=38, n=33), HLA-DR (n=96, n=55). % expression of **B** Ki-67 (n=21) and **C** PD-1 (n=82) expression amongst the three CD4^+^T-cell subsets. **D** Analysis of the four differentiation/cellular states based on KLRG-1 and CD127 expression (n=41). Heatmap shows % expression of each designation, * indicates minimum statistical significance level when compared to both other subsets.

Another hallmark of human T_RM_ is the adoption of a self-restrained, resting state necessary to prevent inflammatory damage to the tissues they reside in[5, 13, 15]. Correspondingly, we observed the highest expression of the inhibitory marker PD-1 on the CD69^HI^ population, and the lowest expression on T-cells lacking CD69 (**Fig.2C**). To investigate cellular activation states in more detail, we also analysed co-expression patterns of killer cell lectin-like receptor-G1 (KLRG-1 – a marker of antigen experience) and CD127, an indicator of common-γ chain cytokine sensitivity, as in human T_RM_ studies[26–28]. CD69^HI^ cells contained the most resting (KLRG-1^-^CD127^+^) cells, whereas CD69^INT^ were enriched for the previously activated (KLRG-1^+^CD127^+^) population (**Fig.2D**).

These data illustrate a phenotypic dichotomy between CD69^HI^ and CD69^INT^ cells, with the former exhibiting features associated with previously described human T_RM_ cells: i) complete exclusion from peripheral circulation, ii) high expression of proteins associated with tissue retention, iii) occupation of restricted tissue niches, and iv) a resting, restrained phenotype. Contrastingly, the CD69^INT^ population had a distinct homing receptor profile and features consistent with recent activation, increased proliferative capacity, and the potential for recirculation.

### Liver CD69^HI^T_RM_ cells and CD69^INT^ cells are skewed towards T_H_1 and T_H_2 functional profiles respectively

CD4^+^T_RM_ cells have a superior functional capacity to circulating T-cells and mediate protection against a number of viral infections in other organs[1, 15]. To assess the functional potential of CD69^HI^ T_RM_ and CD69^INT^ cells, we measured cytokine production upon short-term stimulation. Following T-cell receptor (TCR) ligation with 5-hour anti-CD3/CD28 stimulation, more CD69^HI^ cells responded with IFN-γ expression than any other population, whereas CD69^INT^ cells contained the most IL-4^+^ and IL-2^+^ cells, and more TNF-α^+^ cells than CD69^-^ T-cells (**Fig.3A**).. No differential expression of IL-17 or IL-10 was seen. IFN-γ was also significantly enhanced in CD69^HI^ compared to CD69^-^ cells in both PMA/Ionomycin-stimulated and unstimulated conditions (**Fig.3B, Supp.Fig.4**). Unstimulated CD69^INT^ cells also produced more IL-4 than the CD69^HI^subset.

**Fig.3.**
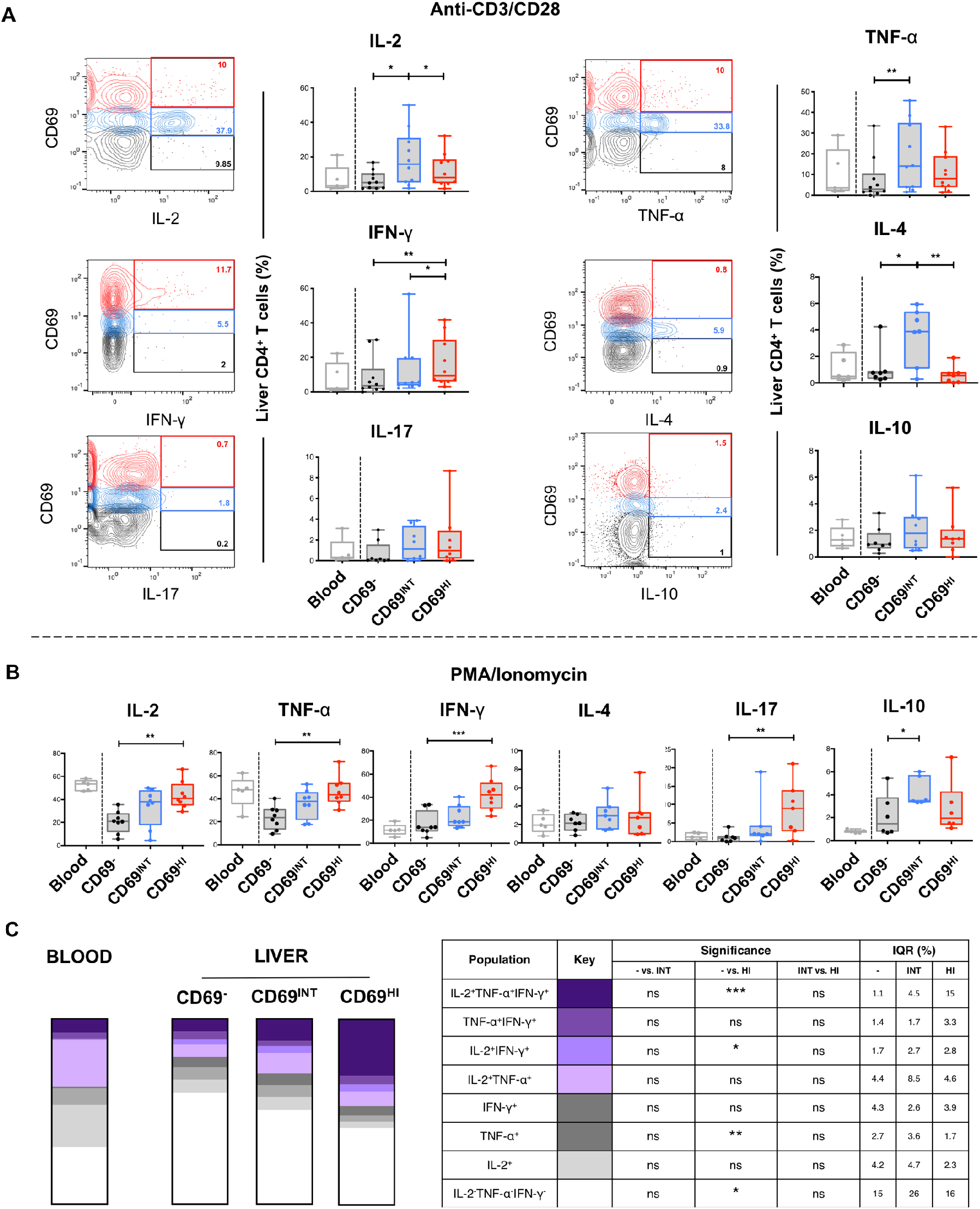
Liver CD69^HI^T_RM_ and CD69^INT^ cells are skewed towards T_H_1 and T_H_2 functional profiles respectively. **A** Intracellular cytokine after 5-hour stimulation of IHL with anti-CD3/CD28. Example flow cytometry plot; gate frequencies indicate % producing cytokine in just the CD69-expressing CD4^+^T-cell subset shown (indicated by gate colour). Reference CD69^-^ PBMC included for reference and not included within statistical comparisons (whiskers represent 5-95% CI). **B** Intracellular cytokine after 5-hour stimulation of IHL with PMA and Ionomycin. **C** Multi-functional responses of blood CD69^-^; and liver CD69^-^, CD69^INT^, and CD69^HI^ CD4^+^ T cells following 5-hour PMA/Ionomycin stimulation. Stacked bar chart heights represent median % of each combination of IL-2/TNF-α/IFN-γ-expressing cells as shown (Liver, n=7; Blood, n=5). Table displays statistical significance between the three liver populations, and interquartile range values.

Human T_RM_ from other organs are often multifunctional, producing the cytokines IFN-γ, TNF-α and IL-2 simultaneously, a property that equips T-cells for better pathogen control[29–33]. Likewise, we observed an increase in type-1 multifunctional cells in CD69^HI^T_RM_ compared to CD69^-^, a feature not shared by CD69^INT^ cells (**Fig.3C**). Thus, CD69^HI^ and CD69^INT^ cells are functionally distinct, with CD69^HI^ favouring more IFN-γ and type-1 multifunctional responses, and CD69^INT^ cells predisposed to enhanced IL-4 production.

### Increased CD69^INT^ frequencies are associated with fibrosis progression in chronic HBV infection

To test if activated CD69^INT^ correlated with disease progression features, we analysed MELD scores (a commonly used metric to assess severity of non-viral chronic liver disease[34]) in explants with chronic hepatitis. There was no correlation with any CD69 subset and MELD score, irrespective of liver disease aetiology (**Supp.Fig.5A**). Stratifying patients into autoimmune and dietary liver disease groups revealed no disease-specific enrichment of any CD4^+^T-cell population (**Supp.Fig.5B**). However, we did observe a modest yet consistent reduction in CD69^HI^ cell frequencies in patients with chronic hepatitis B infection (CHB) (**Fig.4A**).

**Fig.4.**
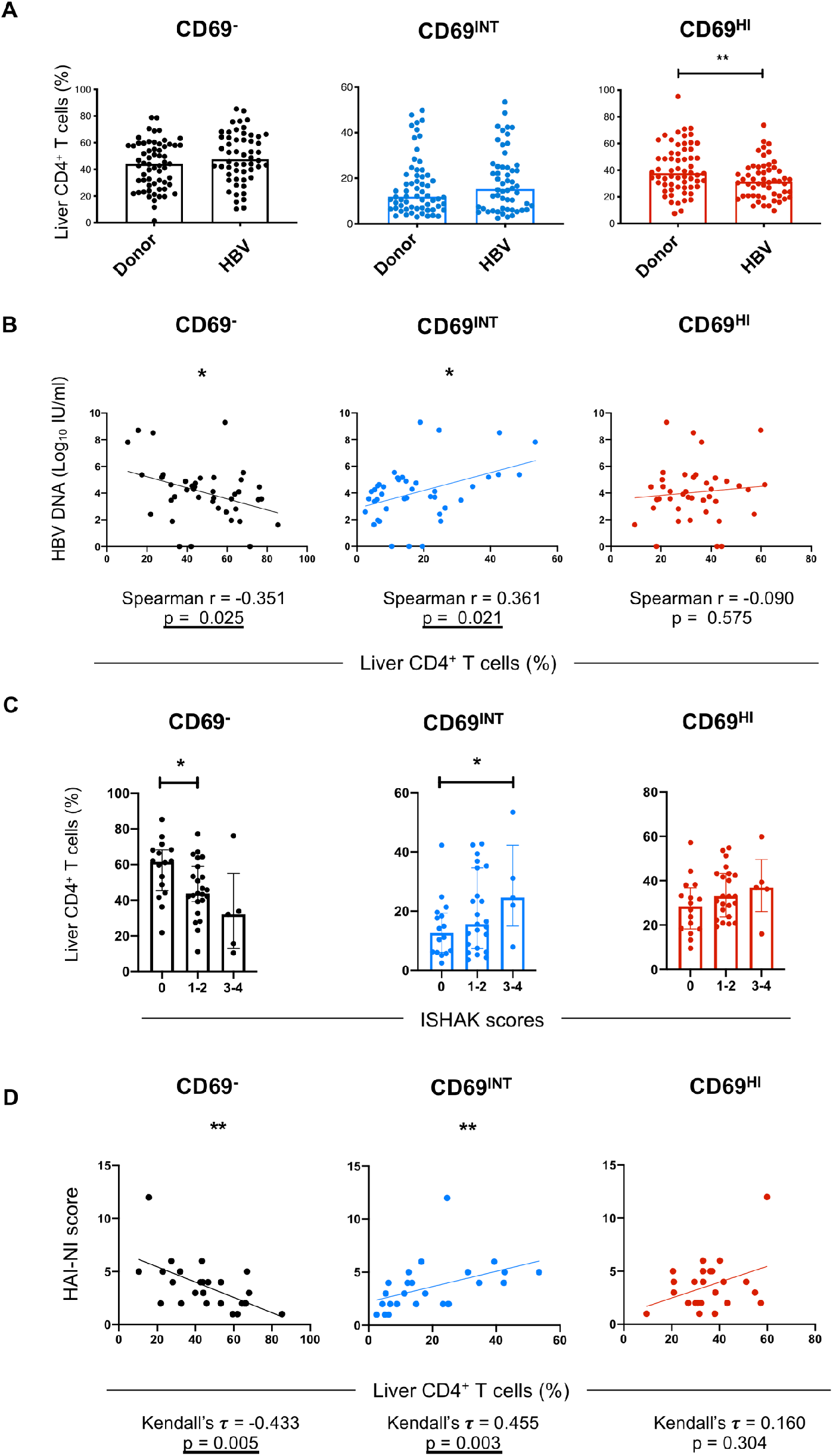
An increase in CD69^INT^ cells is associated with fibrosis progression in CHB infection. **A** Representation of each subset in healthy donor livers (n=62) and patients with chronic HBV (n=54). Healthy donor livers comprised of healthy donor explant transplant rejections(n=11), healthy tissue biopsies (n=5), colorectal cancer margin liver explant (n=36), HCC margin liver explants (n=8) and cyst-free areas from polycystic liver disease explants (n=2). **B** HBV DNA, ISHAK scoring (**C**), and HAI-NI scoring (**D**) plotted against % of each subset in HBV donors. Correlation and p values reported for each plot.

Progression to advanced fibrosis in HBV-infected individuals is highly heterogeneous, with the duration of infection and phase of disease contributing to this process[35]. When analysing patients with CHB by hepatitis B ‘e’ antigen (HBeAg) seropositivity, hepatitis B viremia, or extent of liver inflammation using serum alanine aminotransferase (ALT) concentrations[35], we noted that CD69^INT^ frequencies correlated weakly with serum HBV DNA. The presence of HBeAg or extent of liver inflammation had no impact on CD4^+^T-cell subset distribution (**Fig.4B, Supp.Fig.6A-B**). Combined analysis of these three metrics into the distinct clinical phases also revealed no subset-linked association (**Supp.Fig.6C**).

Robust T_H_2 responses have been linked to the promotion of fibrosis in chronic hepatitis C infection[36, 37], and T_H_2 cytokines IL-4 and IL-13 can activate hepatic stellate cells to produce collagen and promote fibrosis[38, 39]. Therefore to address whether the T H2-biased CD69^INT^ cells within the liver are linked to the degree of fibrosis in CHB patients, we sub-categorised the patients using a using the validated ISHAK and histology activity index-necroinflammatory (HAI-NI) scoring systems[40]. Frequencies of CD69^INT^ cells increased with progressive fibrosis, whereas the CD69^HI^T_RM_ cells remained consistent (**Fig.4C**). CD69^INT^ cells were also significantly more frequent in patients with a higher intrahepatic necroinflammatory score (**Fig.4D**). Together, these data point to a putative role for CD69^INT^ CD4^+^T-cells in fibrosis in patients with CHB.

### CD69^INT^ and CD69^HI^ T_RM_ cells are induced by the liver microenvironment

Next, we sought to determine the origin of these distinct liver CD4^+^T-cell subsets, by deconstructing the contribution of different hepatic cell types *in vitro*. To investigate the role of hepatic epithelia (hepatocytes, biliary epithelia), we first cultured peripheral blood-derived CD4^+^T-cells with different hepatocyte cancer derived cell lines (Huh-7, HepG2, Hep3B). In as little as 16 hours there was robust induction of intermediate-level CD69 expression (**Fig.5A**). By contrast, neither primary hepatic sinusoidal endothelial cells (HSEC), primary biliary epithelial cells (BEC), nor hepatic stellate cell line LX-2 were able to induce intermediate CD69 expression during the same time frame. However, primary BEC were able to strongly induce CD69^INT^ cells from three days onwards, suggestive of a feature of liver epithelia necessary for generation of this phenotype (**Supp.Fig.7A-B**). Hepatic epithelia-induced CD69 upregulation to an intermediate level was not reflective of an activation event, as conventional T-cell activation with anti-CD3/CD28 stimulation led to only high CD69 expression, and no concomitant upregulation of activation marker CD38 was seen (**Fig.5A-B**). Mechanistically, this intermediate CD69 induction required direct T-cell-epithelial cell contact (**Supp.Fig.7C**). *In vitro*-generated CD69^INT^ cells recapitulated the complex phenotypic signature of liver CD69^INT^ cells observed *ex vivo*, showing upregulation of S1PR1, CXCR1, CXCR3 and CX_3_CR1 after just 5-hours in co-culture (**Fig.5C**). CD69^INT^ cells generated by Huh-7 co-culture were also capable of producing IL-4 upon stimulation, resembling intrahepatic CD69^INT^T-cells (**Supp.Fig.8, Fig.3A**). Thus, short-term contact with hepatic epithelia can drive CD69^INT^ cell development *in vitro*.

**Fig.5.**
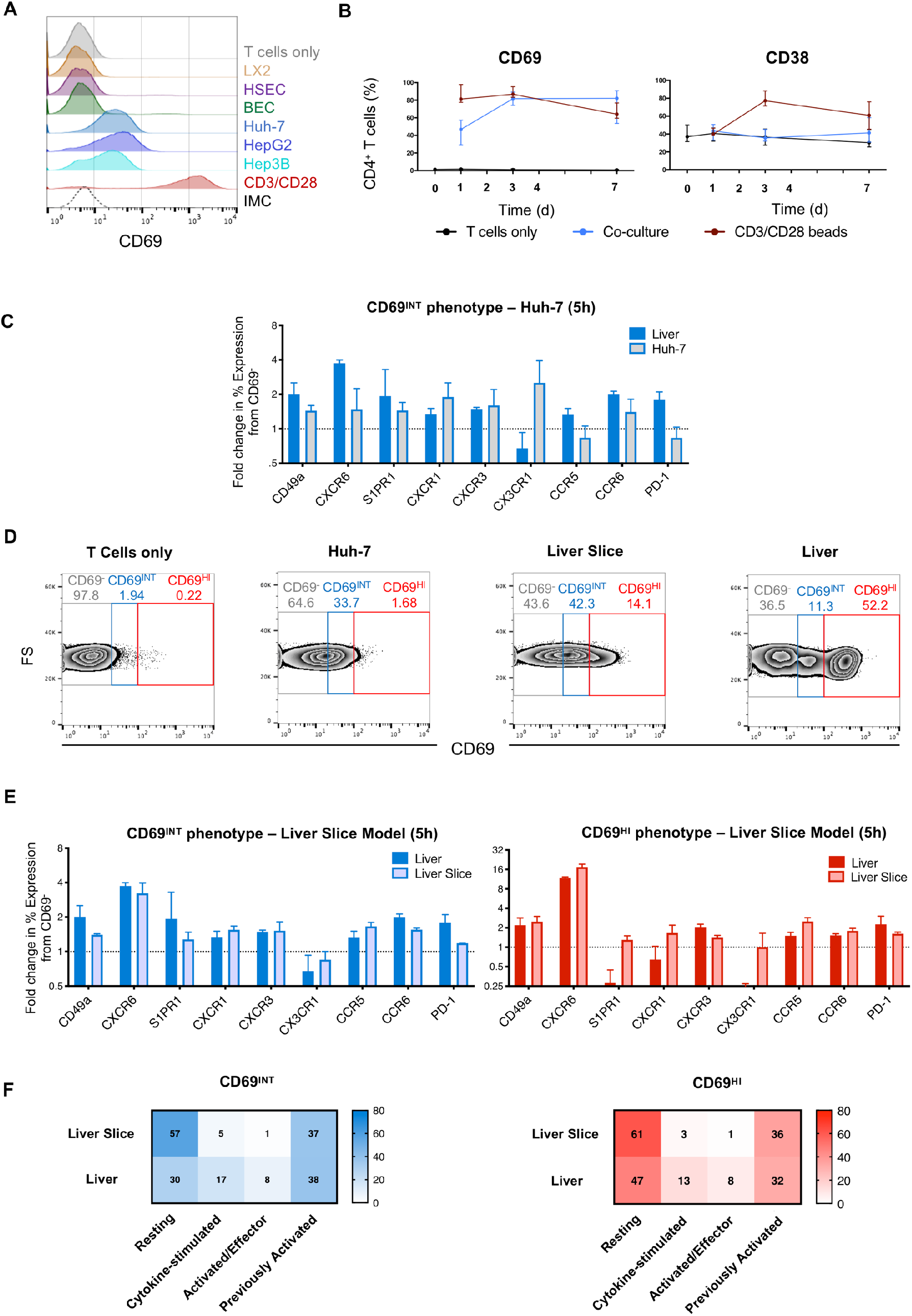
Different components of the hepatic niche can instruct the development of CD69^INT^ and CD69^HI^ T_RM_. **A** % CD69 expression on PBMC-derived CD4^+^T-cells cultured for 16-hours with primary HSEC, primary BEC; hepatic stellate cell line LX-2; hepatocyte cell lines Huh-7, HepG2 or Hep3B; with anti-CD3/CD28; or alone. Histogram displays representative CD69 expression levels in each condition. **B** % CD69 and CD38 expression on blood CD4^+^T-cells over a 7 days culture period with Huh-7 (n=8-10/timepoint). **C** Comparison of key phenotypic markers in Huh-7-generated CD69^INT^ cells from PBMC following 5-hour culture, and matched patient IHL CD69^INT^ cells (n=2). Data in each condition normalised to control CD69^-^ cells in same condition. **D** Representative flow plots showing degree of CD69^INT^ and CD69^HI^ generation within PBMC after 5-hours of culture: alone, with Huh-7 cells, with precision-cut donor-matched liver slices; or from directly isolated IHLs from matched human liver. **E** Comparison of CD69^INT^ cells (left), and CD69^HI^ cells (right) generated from donor-matched PBMCs in precision-cut liver slice model, with matched donor-derived Liver subsets (n=2). Data normalised to condition-matched CD69^-^ cells. **F** Activation/differentiation statuses of CD69^INT^ and CD69^HI^ cells in the different conditions as assessed by KLRG-1/CD127 co-staining patterns (as in **Fig.2D**). Colour intensity and displayed numbers represent median % in each KLRG-1/CD127 designation.

We next asked whether additional signals from the liver microenvironment were required to generate CD69^HI^ cells. To investigate this, we used a co-culture model of patient-derived PBMCs with autologous precision-cut liver slices to allow full retention of the native liver microenvironment (**Fig.5D**). Co-culture of autologous PBMC for 5-hours with matched liver slices led to a modest appearance of both CD69 expressing subsets, not seen with hepatic epithelia co-culture. Remarkably, slice-culturegenerated CD69^HI^ cells phenotypically resembled *ex vivo* intrahepatic CD69^HI^ cells, with high expression of CXCR6, CD49a, CCR5 and PD-1, low expression of S1PR1 and a largely resting (KLRG-1^-^CD127^+^) phenotype (**Fig.5E-F**). Correspondingly, CD69^INT^ cells also generated through hepatic slice culture acquired many of the phenotypic characteristics of their *ex vivo* counterparts, notably expression of CX_3_CR1 and CXCR3, and partial acquisition of the residency markers CD49a and CXCR6. Together these results suggest that CD4^+^T-cell contact with hepatic epithelia promotes their differentiation to a CD69^INT^ phenotype in the liver, whereas CD69^HI^T_RM_ generation requires additional signals present in the liver microenvironment.

## DISCUSSION

In this study, we uncovered two distinct cell types in the human liver – a prototypical CD69^HI^ CD4^+^T_RM_ subset with a tissue-retention signature, and a resting restrained phenotype with the ability to instigate type-1 multifunctional responses on stimulation; and a novel subset of CD69^INT^CD4^+^T-cells with a CXCR3^+^CXCR1^+^CX_3_CR1^+^ phenotype that was more activated, recirculating and skewed towards T_H_2 responses on stimulation. We show that these two cell types reside in distinct niches in the liver, have different generation requirements, and are equipped to play distinct roles in liver disease.

In agreement with other human CD4^+^T_RM_ studies, liver CD69^HI^ cells expressed T_RM_-associated retention molecules CD49a and CXCR6[8, 18, 41], showed low expression of homing receptors S1PR1 and CX_3_CR1[13], a resting and restrained phenotype including high PD-1 expression[13, 41], and the potential to robustly produce T_H_1 cytokines[32, 33, 41]. CXCR6 is of particular importance as a key liver retention molecule that is required for residence of multiple lymphocyte types in the liver[42–44].

We recently described human liver CD8^+^T_RM_ that share some of these key features (CXCR6^+^, PD-1^+^ and rapid functionality)[5]. Intriguingly, intrahepatic CD8^+^T_RM_ in both mouse and humans are thought to reside in a unique niche, with recent studies revealing their presence in the liver vasculature[5, 45, 46]. Our data also suggests that liver CD4^+^T_RM_ can be found within sinusoids. Candidate molecules for maintaining the CD69^HI^T_RM_ in this niche include CXCR6 through interactions with its ligand CXCL16, expressed on the sinusoidal lumen[42, 47], or integrin αLβ2 - ICAM interactions[46]. Furthermore, our findings suggest CD4^+^CD69^HI^T_RM_ can be found at the outer edges of portal aggregates, likely directed specifically to portal vasculature by CCR5 ligands[48]. The strategic positioning of CD4^+^T_RM_ in both these vascular sites could allow efficient immunosurveillance and opportunities to interact with other key immune cells within the liver.

CD69^INT^CD4^+^T-cells have not been previously described. Here we show that they were enriched in the liver in health and disease, but were also present in the blood and other lymphoid and non-lymphoid organs, and retained expression of S1PR1. To our knowledge this is the first report of concomitant CD69 and S1PR1 expression, suggestive of a required threshold of CD69 expression to induce full S1PR1 downregulation[49]. Functionally, CD69^INT^ cells were most able to produce T_H_2 cytokine IL-4. As T_H_2 responses have been consistently linked to liver fibrosis in other settings[36–39], this provides one explanation for the observed correlation between CD69^INT^ frequencies and fibrosis in CHB.

CD69^INT^ cells contained the most activated and proliferative cells, features associated with recent immigrants from the blood[26]. Key distinguishing features of hepatic CD69^INT^ cells were expression of CXCR3 and CXCR1 required for hepatocyte homing[24], and importantly, CX_3_CR1. CD69^INT^ cells were also found in hepatic lymph nodes, consistent with a wide-ranging immune surveillance role, analogous to the CX_3_CR1^INT^ ‘peripheral memory’ CD8^+^T-cells that survey peripheral tissues in both humans and mice[22, 23]. Further, a small population of non-resident CD69^INT^ CD8^+^T-cells in mice has been previously reported[49]. The ‘migratory memory’ CD4^+^T-cells described by Watanabe *et al*. showed variable CD69 positivity and recirculated through the skin slower than conventional T_CM_[50]. These migratory memory T-cells also produced IL-4, similar to liver CD69^INT^ T-cells[50].

Our data revealed insights into the mechanisms behind the generation of both CD69^INT^ and CD69^HI^ cells. CD69^INT^-like cells were generated following short-term direct contact with hepatic epithelial cell lines, and primary BEC. Although the molecular mechanism for this remains undefined, *in situ* hepatocyte-contact may cause CD4^+^T-cell CD69 upregulation to an intermediate level, likely increasing liver dwell time and allowing more efficient immunosurveillance. CD69^HI^ on the other hand required additional signals from the liver microenvironment – as cells with a highly concordant phenotype could be formed from blood-derived CD4^+^T-cells when cultured with autologous liver slices. Cytokines including IL-15 and TGF-β have been shown to drive adoption of the CD8^+^T-cell tissue residency program[11], and we previously demonstrated that combinations of both these cytokines were sufficient to generate cells with a liver CD8^+^T_RM_ phenotype[5]. This raises the possibility that these cytokines provide the same additional signals for CD4^+^CD69^HI^ T_RM_ formation.

In conclusion, this study provides a comprehensive framework for understanding liver CD4^+^T_RM_ biology, and characterises a novel CD69^INT^ population that is linked to fibrosis progression in CHB. We suggest that for at least some peripheral tissues, binary expression of CD69 alone is not sufficient to define resident CD4^+^T-cells. This work will facilitate the understanding of the role of liver CD4^+^T-cells in hepatic immune homeostasis, with implications for the development of novel immunotherapeutic strategies for chronic liver diseases.

## MATERIALS AND METHODS

### Ethical Approval

This study was approved by local research ethics committees in Birmingham and London. All samples were obtained with written informed patient consent. All study protocols complied with the 1975 Declaration of Helsinki.

### Patient Samples and immune cell isolation

Blood, liver and lymph node samples from centre A, the Queen Elizabeth Hospital, Birmingham: reference: 06/Q2702/61 and 06/Q2708/11. Blood, liver (resections, biopsies and fine needle aspirates), gut, spleen and lymph node samples from centre B were obtained from either the Royal Free Hospital, London: references: 16/WA/0289, 11/WA/0077, 11/H0720/4 [RIPCOLT clinical trial number 8191] or 11/LO/0421 or Royal London Hospital, Barts Health NHS Trust: references: P/01/023, 16/LO/1699 or 17/LO0266. Immune cells were isolated from tissues and blood through tissue digestion and density centrifugation (see **Supplementary Experimental Methods**). See **Supp.Table 1** for full patient details.

### Flow cytometry

For surface staining, cells were incubated with fluorescence-conjugated antibodies on ice for 20-30min. For intracellular staining, cells were either fixed with 1% formaldehyde (Sigma-Aldrich) for 15min, permeabilised with 0.1% Saponin (Sigma-Aldrich) and stained with relevant antibodies in 0.1% saponin (30min, 20°C); or fixed and permeabilised with Cytofix/Cytoperm (BD Bioscience) or FoxP3 Buffer Set (BD Bioscience) according to the manufacturer’s instructions, and staining performed in 0.1% Saponin. Dead IHL were identified (and excluded from analysis) using either a fixable live/dead dye (Thermo-Fisher) for all centre B samples, or zombie dyes (Biolegend) for all cultured centre B samples. Samples were analysed on an ADP CyAn flow cytometer running Summit software (Beckmann Coulter; centre A), or LSRII or X20 flow cytometers ruining FACSDiva software (BD Bioscience) for samples from centre B. See **Supp.Table 2** for list of antibodies used and **Supp.Fig.1** for gating strategies.

### Immunofluorescence

Formalin-fixed 3μm liver biopsy sections were deparaffinised with xylene, rehydrated with 99% industrial denatured alcohol, and underwent antigen retrieval by microwaving in Tris-based antigen unmasking solution (Vector Labs). REAL peroxidase Blocking agent (Dako, Agilent) and casein solution (Vector labs) were added sequentially for 30 mins each, before primary antibodies diluted in TBS + 0.1% Tween (TBST) were added for 1h. For antibodies used, see **Supplementary Experimental Procedures**. Following three washes with TBST, secondary antibodies were applied for one hour in TBST, autofluorescence quenched with the TrueVIEW autofluorescence quenching kit (Vector Labs), and tissues mounted with VECTASHIELD^®^ Vibrance™ Antifade Mounting Medium (Vector Labs). Tissues were imaged using the Zeiss LSM 880 microscope (Carl Zeiss Ltd) equipped with a x63 water immersion objective.

### T-cell stimulation for assessment of cytokine production

PBMCs/IHLs were cultured alone, with 1:1 ratio of anti-CD3/CD28 beads (Dynabeads – ThermoFisher) (centre A) or 0.5μg/mL immobilised anti-CD3 and 5μg/mL anti-CD28 (ThermoFisher) (centre B), or 50ng/mL phorbol 12-myristate 13-acetate (PMA) and 1μM Ionomycin (both Sigma Aldrich, UK), all with 10μg/mL Brefeldin A (Sigma Aldrich). For culture and media details see **Supplementary Experimental Procedures**.

### CD4^+^T-cell isolation and cell culture

CD4^+^T-cells were isolated from PBMCs with the EasySep™ human CD4^+^T-cell enrichment kit (StemCell Technologies). T-cells/PBMCs were added to cultured hepatic epithelial cell lines (Huh-7, HepG2, Hep3B), hepatic stellate cell line LX-2, primary hepatic sinusoidal endothelial cells (HSEC), and primary biliary epithelial cells (BEC). Primary BEC and HSEC were isolated inhouse as previously described[19, 20]. For media details see **Supplementary Experimental Procedures**. 1×10^6^ PBMCs/T-cells were added to each well, and cultured for up to 7 days. For transwell separation experiments, T-cells were added to top of 0.4μm pore transwell insert (Corning) separated from hepatic cells at the bottom of the 24-well plate.

### Liver slice cultures

Precision-cut-liver slices were prepared using a TruSlice tissue slicer (CellPath) and cultured in complete DMEM (with 2% FBS) in 48-well plates. Autologous PBMCs were added on top in T-cell media, and plates cultured for 5hr at 37°C before PBMC harvest and use in downstream assays.

### Data analysis and statistics

All flow cytometry data were analysed using FlowJo v.9-10 (FlowJo LLC). Statistically tested was done in Prism v.8 (GraphPad). Median average values were used throughout, and non-parametric testing used throughout as follows: Wilcoxon matched-pairs signed rank tests for comparison of two paired groups, Freidman tests with Dunn’s multiple for testing 3+ paired groups, Mann-Whitney tests for two unpaired groups, Kruskall-Wallis tests with Dunn’s multiple post-hoc tests for testing 3+ unpaired groups, Spearman’s Rank. Order Correlation for comparing two continuous variables, and Kendall’s Tau rank correlation tests for comparing frequencies with ranked scoring systems. Significance levels defined as *p<0.05, **p<0.01, ***p<0.001, ****p<0.0001. Liver images in graphical abstract and **Fig.5D** reproduced from Servier medical art (Les Laboratoires Servier) under a creative commons licence (https://smart.servier.com/).

## List of abbreviations

ALD: alcoholic liver disease
ALT: alanine aminotransferase
BEC: biliary epithelial cells
CHB: chronic hepatitis B virus
CTLA-4: cytotoxic T-lymphocyte-associated protein-4
FNA: fine needle aspirates
HAI-NI: histology activity index necroinflammation score
HBeAg: hepatitis B e antigen
HBV: hepatitis B virus
HCC: hepatocellular carcinoma
HSEC: hepatic sinusoidal endothelial cells
IFN-γ: interferon-gamma
IHL: intrahepatic leukocytes
IL: interleukin
KLRG-1: killer cell lectin-like receptor-G1
MFI: median fluorescence intensity
NASH: non-alcoholic steatohepatitis
PBC: primary biliary cholangitis
PD-1: programmed cell death protein-1;
PSC: primary sclerosing cholangitis
S1PR1: sphingosine-1-phosphate receptor-1
T_CM_: central memory T cell
TCR: T cell receptor
T_EM_: effector memory T cell
TGFβ: transforming growth factor beta
T_N_: naïve T cell
TNFα: tumour necrosis factor-alpha
T_REG_: regulatory T cell
T_RM_: tissue-resident memory T cell.

## Author contributions

BGW, LJP, MKM and ZS conceived the project and designed experiments; BGW, LJP, XL, SPD, OEA, USG, AP, KA, YL and GR generated data, BGW, LJP, XL and SPD analysed data; USG, GR, GH and PTFK provided access to patient material; BGW wrote the manuscript and carried out statistical comparisons; BGW, LJP, SPD and ZS constructed figures; BGW, LJP, MKM and ZS critically evaluated the data and edited the manuscript; YH, MKM and ZS acquired funding and supervised the project, all authors reviewed the manuscript.

## Acknowledgements

The authors are grateful to all patients and donors at the Queen Elizabeth Hospital Birmingham, and the Royal Free Hospital, London. We would like to thank the whole liver transplantation unit and all the clinical pathology team at the Queen Elizabeth Hospital for sample allocation; and all clinical staff who helped with patient recruitment across both centres including the Tissue Access for Patient Benefit project at The Royal Free Hospital. Our thanks also to Loraine Brown and Bridget Gunson for their help with access and management of patient information and to the support staff at the UCL Infection and Immunity Flow Cytometry Core Facility.

All work was performed at the Centre for Liver and Gastrointestinal Research, Institute of Immunology and Immunotherapy, University of Birmingham; and within the Division of Infection and Immunity, UCL. All samples collected from NIHR Birmingham Biomedical Research Centre, Royal Free Hospital, London, and the Royal London Hospital, London.

