## Supplementary Information for "The human liver microenvironment shapes the homing and function of CD4^+^T-cell populations"

### Supplementary Experimental Procedures

#### Immune cell isolation

Peripheral blood mononuclear cells (PBMCs) were isolated from human blood by density centrifugation (Lympholyte-H [Cederlane] – centre A, Ficoll-Hypaque plus [GEHealthcare] – centre B). Intrahepatic lymphocytes (IHL) were isolated through either chopping, multiple PBS washes, digestion with a stomacher machine (Seward), filtering out debris and density centrifugation (centre A); or for centre B biopsies – mechanical disruption with cell scrapers, filtration to remove debris, centrifugation. Larger centre B resections and explants: enzymatic digestion (30min incubation at 37°C with 0.02% Collagenase IV [Thermo-Fisher] 0.002% DNase I [Sigma-Aldrich]), mechanical disruption through a GentleMACS dissociator (Miltenyi Biotec), then 70μm filtering and centrifugation as above. Lymph node and spleen mononuclear cells were isolated through manual dissection, GentleMACS dissociation, filtration, and density centrifugation on Lympholyte-H (centre A); or manual dissection, filtration, and centrifugation on a Pancoll (PanBiotech) gradient. Intraepithelial lymphocytes (IEL) were isolated from the duodenum and Ileum of the gut by enzymatic digestion (1hr incubation at 37°C shaking [180rpm] with 0.15% Collagenase IV [Thermo-Fisher] 0.02% DNase I [Sigma-Aldrich] 0.0025% Hyaluronidase Type IV-S [Sigma-Aldrich] 0.00125% Liberase DL [Roche]), mechanical disruption using a syringe, then 70μm filtration and centrifugation on a Pancoll gradient as above. The isolation of IHLs from FNA samples was undertaken as previously described[35]. In brief, samples were centrifuged; the remaining cell pellet was re-suspended in 2mL of red blood cell lysis buffer (Biolegend) for 5min on ice, prior to staining. All samples were used immediately. Any samples for later use were frozen in 10% DMSO (Sigma-Aldrich) in fetal bovine serum (FBS) and stored in accordance with the Human Tissue Act.

#### Antibodies for Immunofluorescence

The following antibodies were used: mouse anti-human CD4 (Novus Biologicals; NBP2-46149), mouse anti-human NKp46 (R&D systems, UK; MAB1850), Rabbit anti-human CX3CR1 (Thermofisher, UK; 702321) and rabbit-human CXCR6 (Abcam, UK; ab8023). Single stains and IMCs were used to ensure specificity. Secondary antibodies were all purchased from Thermo-Fisher Scientific.

#### T cell culture for stimulation experiments

PBMCs/IHLs were pre-stained with antibodies against CD69, CD4, CD56, and γδ-TCR (centre A only), washed twice in PBS, and plated out in 96-well plates at $10^6$ cells/well in T-cell media (RPMI [ThermoFisher] + 10% FBS (Sigma-Aldrich), 100U/ml penicillin, 100μg/ml streptomycin, 1% non-essential amino acids (NEAA), 1% L-glutamine [all ThermoFisher Scientific, UK]) containing T cell stimulants.

#### T cell co-culture experiments

Hepatic cell lines all cultured in 24-well plates in 1ml media as follows: Huh-7, HepG2, Hep3B – all in complete DMEM (ThermoFisher): DMEM + 10% FBS, 100U/ml penicillin, 100μg/ml streptomycin, 1% NEAA, 1% L-glutamine; LX-2 – as above but 2% FBS; primary BEC – 1:1 Ham's F12 media (ThermoFisher) & DMEM + 10% heat activated human serum (HD supplies) 2mM Penicillin/Streptomycin, 10μg/L epidermal growth factor, 10μg/L hepatocyte growth factor (both Peprotech), 124 IU/L Insulin, 20μg/L Hydrocortisone (both QE hospital pharmacy, Birmingham), 10μg/L Cholera Toxin, 0.2nM Tri-iodothyronine (both Sigma-Aldrich); primary HSEC – Human Endothelial Serum free media (ThermoFisher) + 10% heat activated human serum, 10μg/L hepatocyte growth factor, 10μg/L vascular endothelial cell growth factor (Peprotech). BEC and HSEC cultured on type-1 rat-tail collagen (Sigma Aldrich, UK) coated plates (coated with 40μg/ml solution). Once adhered to plate, T cells added to 1ml wells in 100μl of T-cell media.

**Supplementary Figures:**

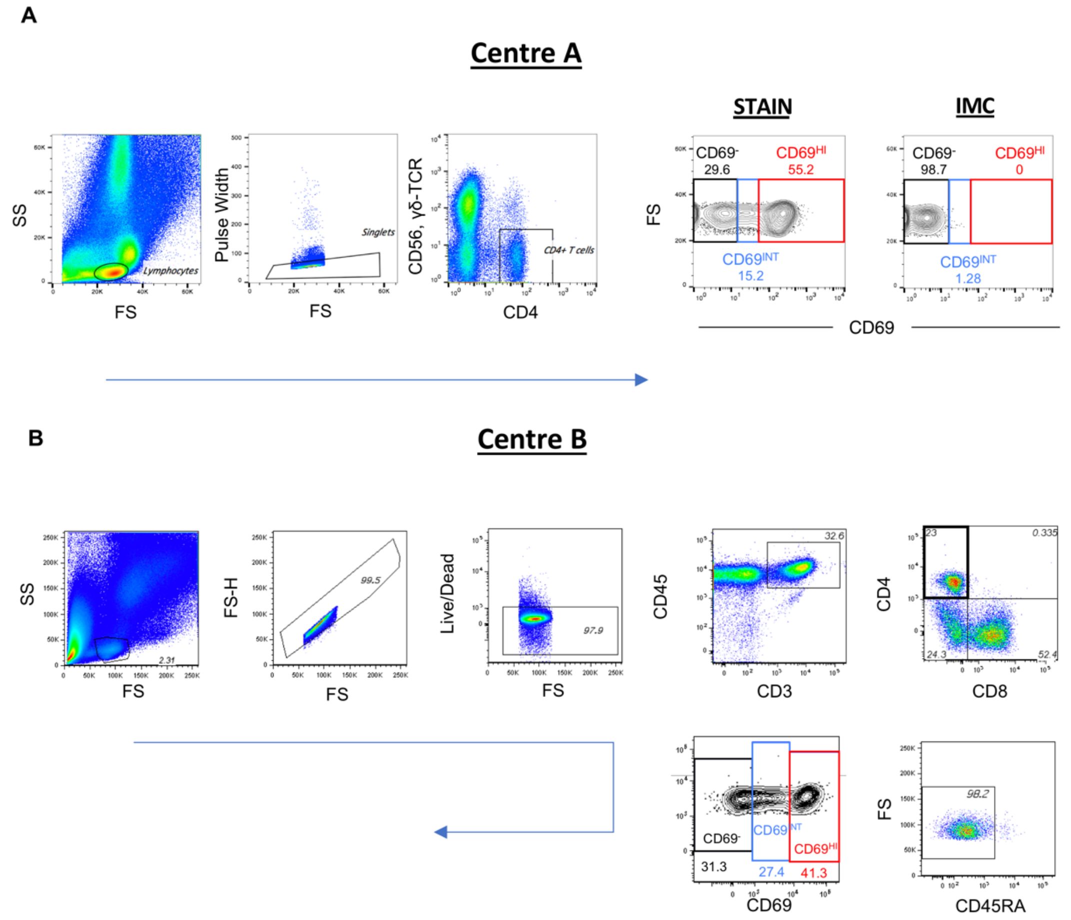


**Supplementary Figure 1 – Gating strategies used to identify and compare hepatic CD69$^-$, CD69$^{INT}$ and CD69$^{HI}$ cells.** Gating strategy used in centre A (**A**), and B (**B**) where blue arrows represent direction of gating (onward gate in bold when multiple in one plot).

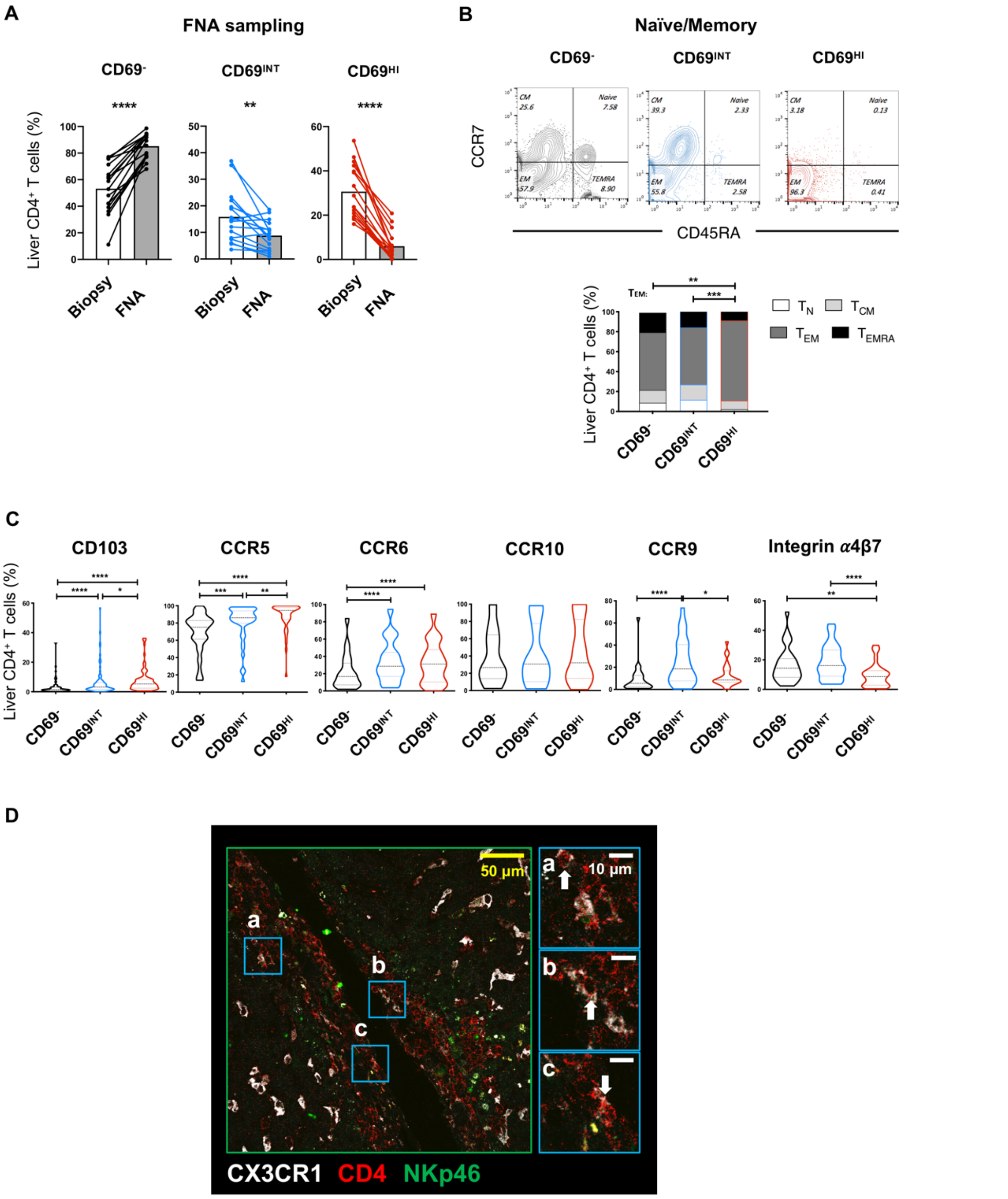


**Supplementary Figure 2 – Homing, location, and naïve/memory profiles of CD69-deliniated subsets. A –** Comparison of each subset (%) recovered from liver tissue digests or fine needle aspirate (FNA) samples. **B** – proportion of liver CD4$^+$ T cells in each subset that comprise naïve, central memory ($T_{CM}$), effector memory ($T_{EM}$), or $T_{EMRA}$ as shown in representative plots and combined stack chart (n=51). **C –** Expression of additional homing/retention molecules in each subset (CD103 n=89, CCR5 n=40, CCR6 n=42, CCR10 n=10, CCR9 n=27, Integrin $\alpha4\beta7$ n=31). **D -** Immunofluorescent staining of formaldehyde-fixed paraffin-embedded liver sections (from patient with PBC). White arrows indicate CX$_3$CR1$^+$ CD4$^+$ T cells.

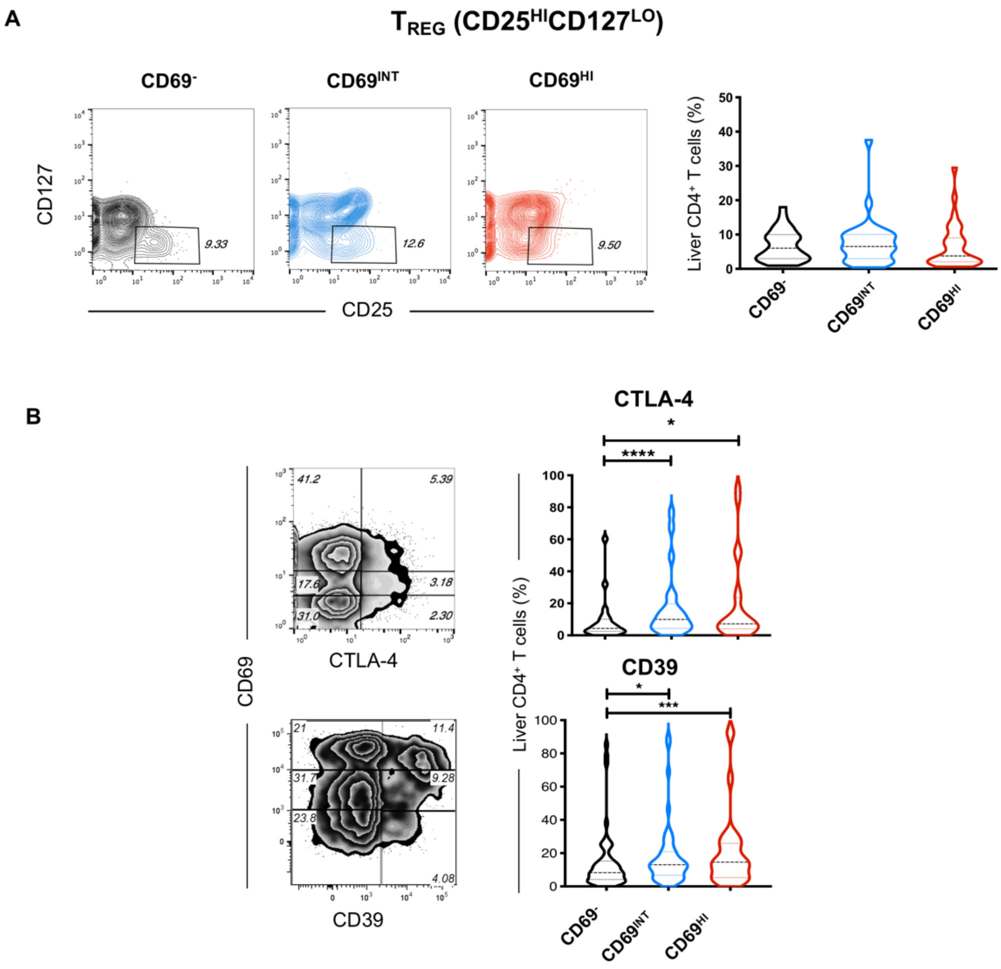


**Supplementary Figure 3 – Conventional regulatory T cells are not enriched in any CD69-designated subset. A** – Representative gating for $T_{REGS}$ ($CD25^{HI}CD127^{LO}$) in each subset. Violin plot shows combined expression data (n=31). **B** – Expression of $T_{REG}$ cardinal features in each subset – CTLA-4 (n=22) and CD39 (n=52) by representative staining and combined total data.

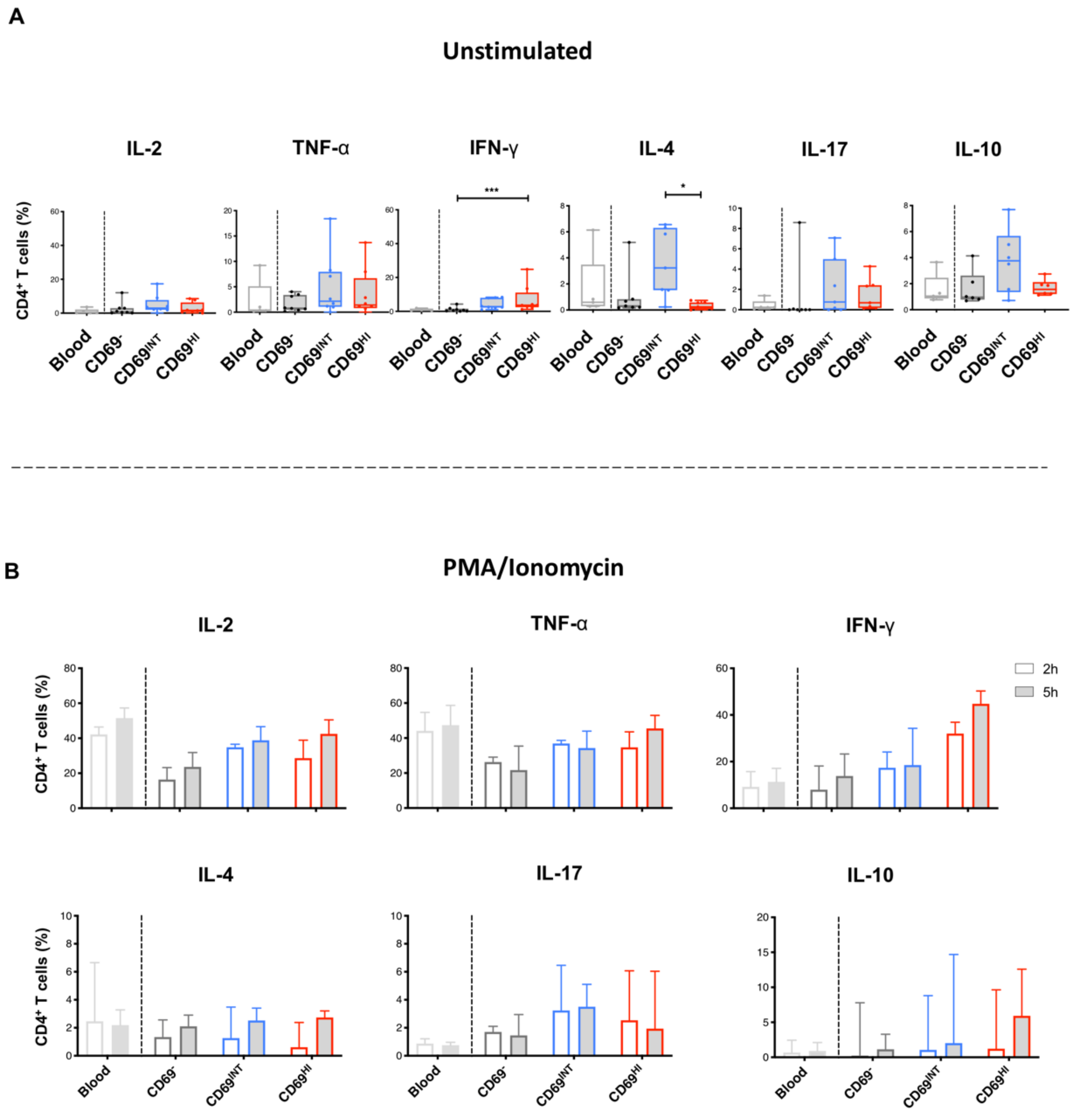


**Supplementary Figure 4 – Differential cytokine responses of intrahepatic subsets marked by CD69 expression.** As in Figure 3, combined intracellular cytokine staining of IHL/PBMCs, but following 5 hours in culture in the absence of stimuli.

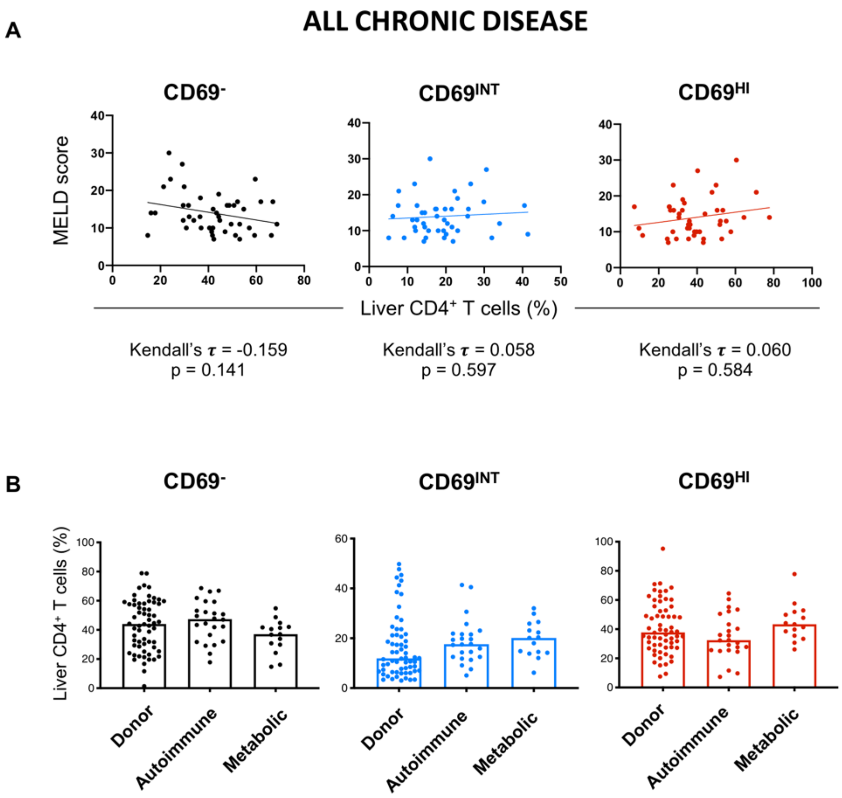


**Supplementary Figure 5 – CD69$^{INT}$ and CD69$^{HI}$ frequencies in non-viral chronic liver diseases. A –** Correlation analysis of patient MELD scores vs. % of each of the 3 subsets for all donors with end-stage liver disease from centre A. **B –** Comparison of subset frequencies in Donor (n=61, composition as in Fig. 5), autoimmune liver disease (n=15 [6 PBC, 8 PSC, 1 AIH]), and dietary liver disease (n=24 [16 ALD, 8 NASH]).

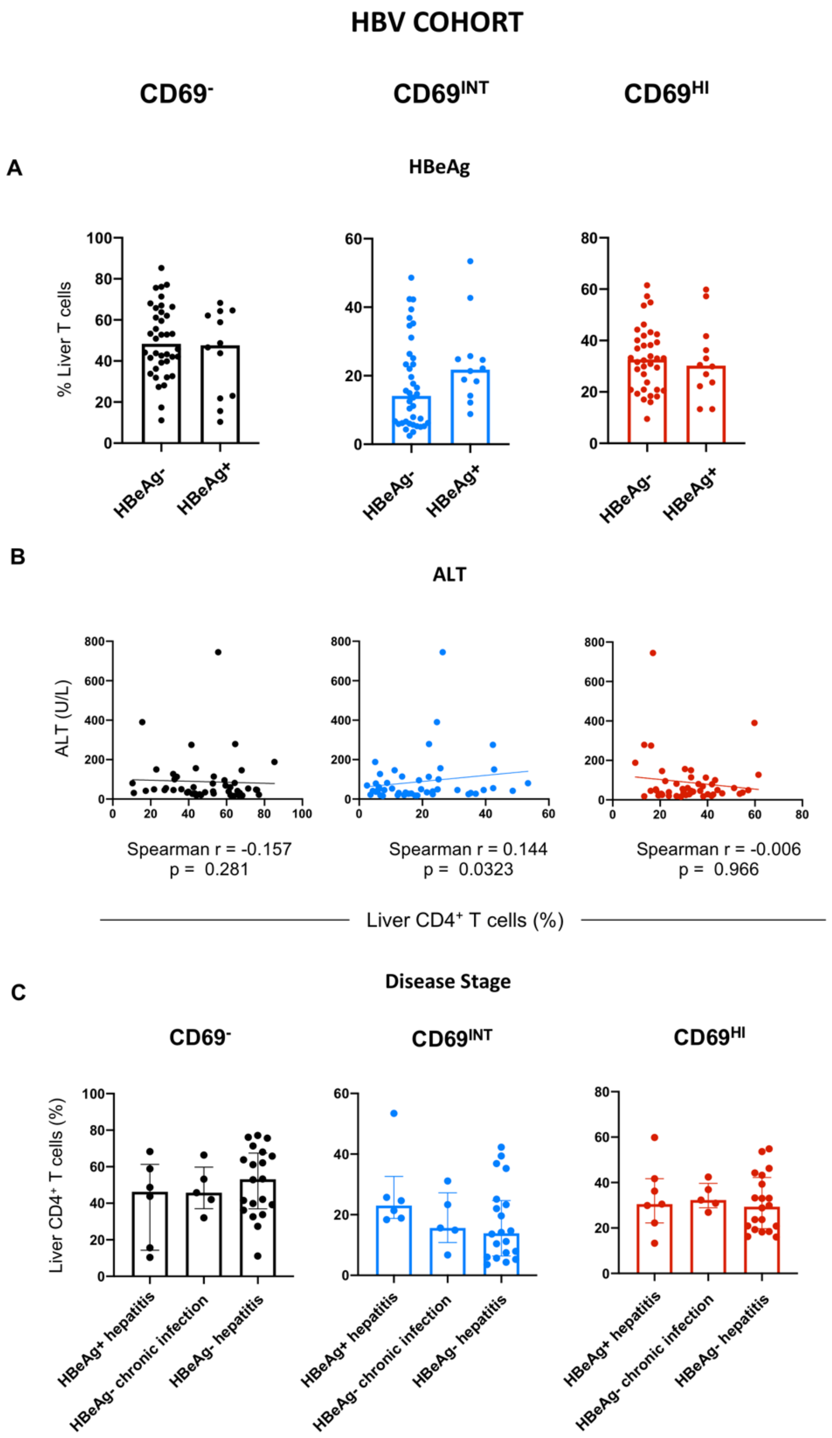


**Supplementary Figure 6 – Additional correlation analysis with clinical parameters in HBV patient livers. A –** Frequency of each subset in Hepatitis B e antigen positive vs – negative patients. **B –** Correlation of patient serum ALT against frequency of each subset. Spearman's correlation analysis p and r values given. **C –** Frequency of each subset in chronic HBV patient donor livers at each HBV disease stage. Disease staging of HBV donors determined by combination of HBeAg, HBV DNA, and serum ALT. Only 1 donor was at immunotolerant stage, so this stage excluded from dataset shown.

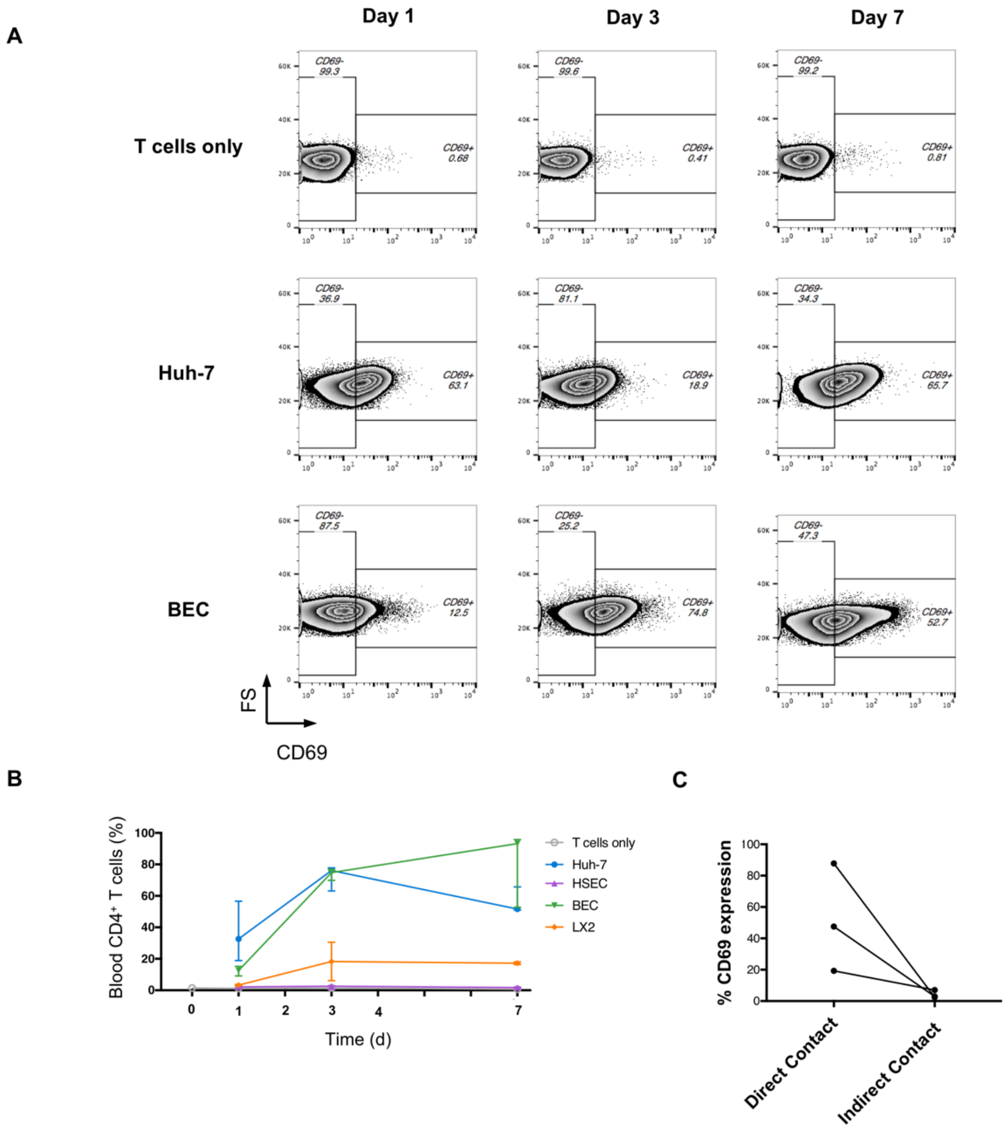


**Supplementary Figure 7 – CD69$^{INT}$ generation occurs with primary human epithelia and is contact-dependent.** CD69 expression in peripheral blood-derived CD4$^{+}$ T cells following co-culture for 1,3, and 7 days alone, with Huh-7 cells, or BEC. Data displayed as representative flow cytometry plots (**A**), and combined donor data alongside HSEC and LX-2 culture conditions (**B**)**.** Median + 95% CI shown (n=3). **C –** CD69% expression amongst PBMC-derived CD4$^{+}$ T cells when either cultured overnight with Huh-7 cells directly (direct contact), or when separated by a 0.4μm pore transwell insert (indirect contact).

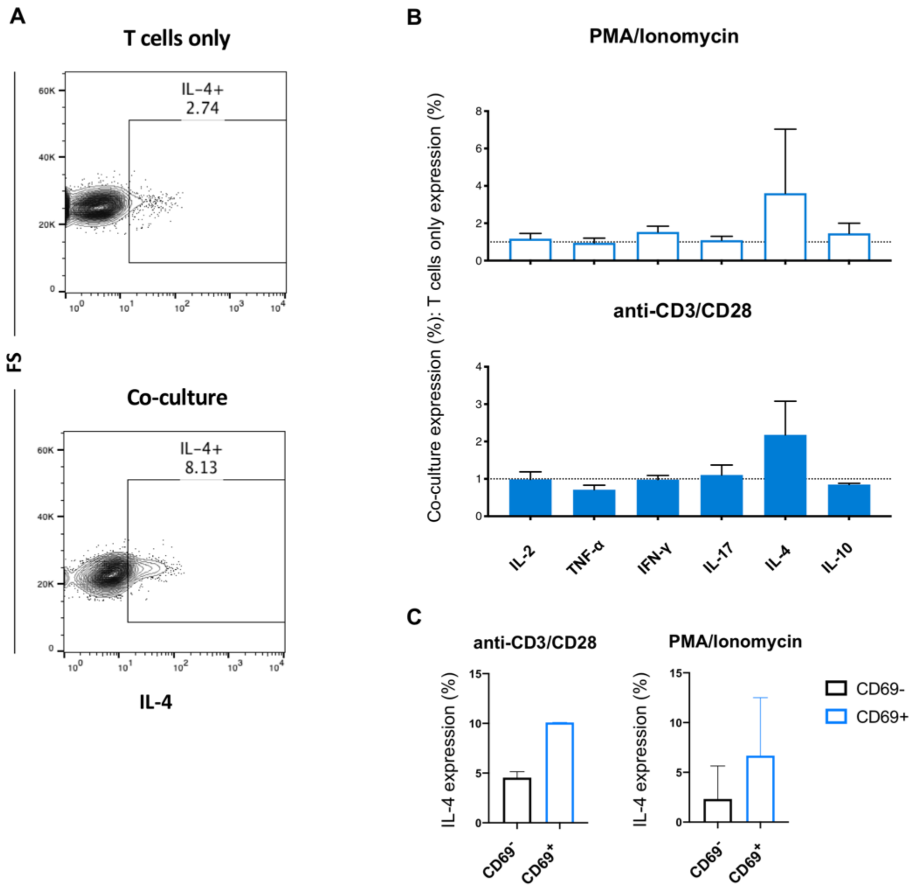


**Supplementary Figure 8 – Co-culture with hepatic epithelia infers CD4+ T cells with IL-4 production capacity.** CD4+ T cells isolated from healthy human blood were cultured for 5 hours either alone or with Huh-7 hepatic epithelial cells, harvested and stimulated for 4 hours with anti-CD3/CD28 or PMA/Ionomycin, and then assessed for their ability to produce prototypic T cell cytokines by intracellular staining and flow cytometry. **A –** Staining example for IL-4 in T cells alone and following co-culture (PMA/Ionomycin stimulation). **B –** % expression of all the cytokines studied in co-cultured cells as a fold change from control (T cell only) cells (anti-CD3/CD28 – top, PMA/Ionomycin stimulation – bottom). **C –** Percentage expression of IL-4 in co-cultured CD69+ versus co-cultured CD69- cells (both stimulation methods shown). Data from this experiment compiled from three independent donors.

### Supplementary Tables:

**Supplementary Table 1 – Patient information for Liver tissue donors.**
Shown are the disease breakdowns, sample type acquired and donor numbers in each group, together with median and IQR of the ages, and gender in each group. Numbers of donors acquired from each centre (A – University of Birmingham, B – University College London). * - 2 patient ages missing, ** - 5 patient ages missing, † - 2 patient genders missing, †† - 4 patient genders missing, ††† - 6 patient genders missing from available data.

| Aetiology | Sample Type | Numbers | Median age (IQR) | % Female | Centre A/B |
|---|---|---|---|---|---|
| Healthy | Explant | 12 | 51 (22)* | 25†† | A(8) B(4) |
| Healthy | Pre-implant biopsy | 5 | 39 (14) | 33† | B |
| Healthy | CRC margin | 36 | 61 (18.5)** | 22†† | B |
| Healthy | HCC margin | 8 | 62.5 (8.8)* | 33† | B |
| Healthy | PLD | 3 | 61 (7) | 100 | A |
| HBV | Explant | 4 | 39 (3)* | 50† | A(1) B(3) |
| HBV | Biopsy | 50 | 35 (17.3) | 30††† | B |
| ALD | Explant | 20 | 59 (14.3) | 15 | A |
| NASH | Explant | 8 | 60.5 (9.3) | 50 | A |
| PBC | Explant | 7 | 44 (11.5) | 71 | A |
| PSC | Explant | 12 | 35.5 (12.3) | 25 | A |
| AIH | Explant | 1 | 21 | 0 | A |
| HCV | Explant | 2 | 59 (6) | 0 | A |
| Cryptogenic | Explant | 2 | 53.5 (10.5) | 50 | A |
| Budd Chiari | Explant | 1 | 31 | 0 | A |
| NCPH | Explant | 1 | 63 | 0 | A |
| SBC | Explant | 1 | 58 | 100 | A |

### Supplementary Table 2 – Antibodies used in flow cytometry experiments

| Antigen | Fluorochrome | Manufacturer | Clone | Catalogue Number |
|---|---|---|---|---|
| **Phenotype** | | | | |
| CCR10 | PE | Biolegend | 6588-5 | 341504 |
| CCR5 | PE | Biolegend | J418F1 | 359106 |
| CCR6 | AlexaFluor488 | Biolegend | G034E3 | 353414 |
| CCR7 | PE-Cy7 | Biolegend | G043H7 | 353226 |
| CCR9 | PerCP-Cy5.5 | Biolegend | L053E8 | 358906 |
| CD103 | APC | BD Biosciences | Ber-ACT8 | 563883 |
| CD103 | BV605 | Biolegend | Ber-ACT8 | 350218 |
| CD103 | FITC | Biolegend | Ber-ACT8 | 350203 |
| CD127 | BV510 | Biolegend | A019D5 | 351332 |
| CD25 | PE-Cy5 | Biolegend | BC96 | 302608 |
| CD27 | FITC | Biolegend | M-T271 | 356404 |
| CD3 | BV711 | Biolegend | OKT3 | 317328 |
| CD38 | APC-Vio770 | Mltenyi Biotec | REA572 | 130-099-151 |
| CD39 | BV421 | Biolegend | A1 | 328214 |
| CD4 | APC | BD Biosciences | RPA-T4 | 555349 |
| CD4 | BV510 | Biolegend | SK3 | 344634 |
| CD4 | FITC | Biolegend | OKT4 | 317408 |
| CD4 | APC-Cy7 | BD Biosciences | RPA-T4 | 557871 |
| CD4 | BV421 | BD Biosciences | RPA-T4 | 562425 |
| CD45 | BUV805 | BD Biosciences | HI30 | 564914 |
| CD45RA | BV421 | BD Biosciences | HI100 | 562885 |
| CD45RA | PE-Cy7 | Biolegend | HI100 | 3014126 |
| CD45RA | eFluor450 | eBioscience | HI100 | 48-0458-41 |
| CD49a | PE | Biolegend | TS2/7 | 328304 |
| CD49d | BV421 | Biolegend | 9F10 | 304322 |
| CD56 | APC-Vio770 | Mltenyi Biotec | REA196 | 130-100-690 |
| CD69 | FITC | BD Biosciences | FN50 | 560969 |
| CD69 | PE-Dazzle594 | Biolegend | FN50 | 310942 |
| CD69 | BV605 | Biolegend | FN50 | 310937 |
| CD8 | PE-Cy5 | Biolegend | RPA-T8 | 301010 |
| CD8a | AlexaFluor700 | eBioscience | OKT8 | 56-0086-82 |
| CTLA-4 | PE-Dazzle594 | Biolegend | L3D10 | 349922 |
| CX3CR1 | PE-Cy7 | Biolegend | 2A9-1 | 341612 |
| CXCR1 | PE-Cy7 | Biolegend | 8F1/CXCR1 | 320620 |
| CXCR3 | AlexaFluor488 | Biolegend | G025H7 | 353710 |
| CXCR6 | APC | Biolegend | K041E5 | 356005 |
| CXCR6 | PerCP-Cy5.5 | Biolegend | K041E5 | 356010 |
| HLA-DR | FITC | BD Biosciences | G46-6 | 555811 |
| HLA-DR | BV421 | Biolegend | L243 | 307636 |
| HLA-DR | Horizon V500 | BD Biosciences | G46-6 | 561224 |
| Integrin β7 | PE | Biolegend | FIB504 | 321204 |
| KLRG-1 | PE | Biolegend | SA231A2 | 367712 |
| PD-1 | PE | Biolegend | EH12.2H7 | 329906 |
| S1PR1 | eFluor660 | eBioscience | SW4GYPP | 50-3639-41 |
| γδ-TCR | APC-Vio770 | Mltenyi Biotec | 11F2 | 130-109-360 |
| **Function** | | | | |
| IFN-γ | APC | BD Biosciences | B27 | 554702 |
| IFN-γ | Horizon V450 | BD Biosciences | B27 | 560371 |
| IL-10 | BV421 | Biolegend | JES3-9D7 | 501421 |
| IL-10 | PE | Biolegend | JES3-9D7 | 501404 |
| IL-17A | PerCP-Cy5.5 | Biolegend | BL168 | 512313 |
| IL-2 | PE | eBioscience | MQ1-17H12 | 12-7029-82 |
| IL-2 | PerCP-eFluor710 | eBioscience | MQ1-17H12 | 46-70290-42 |
| IL-4 | PE-Cy7 | Biolegend | MP4-25D2 | 500824 |
| Ki-67 | PB | Biolegend | Ki-67 | 350512 |
| TGF-β | APC | Novus Biologicals | 1D11 | IC420A |
| TNF-α | eFluor450 | eBioscience | MAb11 | 48-7349-42 |
| TNF-α | FITC | BD Biosciences | MAb11 | 554512 |
